# Different cellular and molecular mechanisms of chitin deposition contribute to the specificity of the two chitin synthases in *D. melanogaster*

**DOI:** 10.1101/2024.12.02.626067

**Authors:** Joan Bertran-Mas, Ettore De Giorgio, Nicolás Martín, Marta Llimargas

## Abstract

Chitin is a major component of arthropod extracellular matrices, including the exoskeleton and the midgut peritrophic matrix. It plays a key role in the development, growth and viability of insects. Besides the biological importance of this aminopolysaccharide, chitin also receives a lot of attention for its practical applications in medicine and biotechnology as a superior biopolymer with excellent physicochemical and mechanical properties. Chitin is produced and deposited extracellularly by chitin synthases. Most insects encode two types of chitin synthases, presumably with type A being required for exoskeletons and type B to produce the peritrophic matrix. It is not fully understood, however, which factors contribute to the specificity of each type of chitin synthase. Here we leverage the advantages of *Drosophila melanogaster* for functional manipulations to evaluate the mechanisms of activity and functional requirements of Kkv (Chitin synthase A) and Chs2 (Chitin synthase B). We first demonstrate that Chs2 is expressed and required in a region in the larval proventriculus that produces the chitin deposited in the peritrophic matrix. We then analyse whether the two chitin synthases can replace each other. We also investigate the subcellular localisation of these chitin synthases in different tissues and their ability to deposit chitin in combination with known auxiliary proteins. Our results indicate that the two different chitin synthases are not functionally equivalent, and that they use specific cellular and molecular mechanisms to deposit chitin. We suggest that the specificity of the different insect chitin synthases may underly the production of chitin polymers with different properties, conferring different physiological activities to the extracellular matrices.

## Introduction

Chitin is an abundant polymer in arthropods and represents a principal component of the apical extracellular matrix, which provides structural and mechanical strength and protects against dehydration, injuries and pathogens. It is found in the exoskeletons, tracheal and internal tendon cuticles, and in the peritrophic matrix (PM). Chitin is composed of linear polymers of β-(1-4)-linked N-acetylglucosamines (GlcNAc) that organise in microfibrils. The chitin chains in the microfibrils can adopt parallel or antiparallel orientations, giving rise to different allomorphs known as α-chitin (antiparallel), β-chitin (parallel) and γ-chitin (parallel and antiparallel). α-chitin forms a highly ordered crystalline structure, while the β- and γ-chitin allomorphs are less tightly packed, more flexible and hydrated (Liu et al., 2019b, Moussian, 2019, Yu et al., 2024, Zhao, 2019, Zhu et al., 2016). Besides its physiological activities, chitin and its deacetylated form, chitosan, are considered most promising biopolymers and serve multiple applications in biotechnology, biomedicine, agriculture, food industry or cosmetics (Behr and Ganesan, 2022, Casadidio et al., 2019, Elieh-Ali-Komi and Hamblin, 2016). Thus, there is a growing interest in understanding the mechanisms of chitin synthesis and deposition to leverage the excellent properties of this biomaterial (Naqvi and Moerschbacher, 2017).

Chitin is synthesised by a dedicated type of enzymes known as chitin synthases (CHS) that belong to the β-glycosyltransferase family, which polymerise and extrude the polymer to the extracellular space (Merzendorfer, 2011). Insects typically encode two different types of CHS, which belong to the class-A and class-B. It is assumed that class-A are expressed and produce chitin in ectodermal tissues while class-B are expressed and synthesise chitin in the gut PM (Liu et al., 2019a, Liu et al., 2019b, Yu et al., 2024, Zhao, 2019, Zhu et al., 2016). The PM consists of a layer of chitin and glycoproteins that lines the midgut and is required for digestion and protection against pathogens, abrasion and toxins. Depending on the insect order, the PM is synthesised as delaminating lamellae along the length of the midgut epithelium (Type I PM), or in a specialised anterior region of the midgut, the cardia (Type II PM) from where it is secreted into the midgut (Hegedus et al., 2009, Hegedus et al., 2019, Merzendorfer, 2016).

Knock-down studies in different insect species have identified specific functional requirements for class-A and class-B CHS, requirements that may respond to their specific pattern of expression in ectoderm or gut, respectively (reviewed in (Zhao, 2019, Yu et al., 2024)). In agreement with this, in *Tribolium castaneum*, CHS1 (type A) RNAinterference affected molting and decreased chitin content in the whole-body, whereas CHS2 (type B) RNAinterference affected feeding and larval size and specifically decreased chitin content in the midgut (Arakane et al., 2005). However, the previous studies have not addressed whether the two different types of insect CHS are functionally equivalent, and whether besides their expression pattern, other factors contribute to their specificity. Here we take advantage of *D. melanogaster* as an ideal model for functional analysis and ask about the specificity and mechanisms of activity of insect class-A and class-B CHS.

*D. melanogaster* encodes *krotzkopf verkehrt (kkv*) and *Chitin Synthase 2* (*Chs2*), which belong to the class-A and class-B CHS, respectively (Gagou et al., 2002, Tellam et al., 2000). Kkv has been largely studied and there is currently extensive information about the functional requirements, the pattern of localisation and its molecular mechanism of activity (Adler, 2019, De Giorgio et al., 2023, Moussian et al., 2015, Moussian et al., 2005, Ostrowski et al., 2002). Kkv is expressed and required for chitin deposition in the ectoderm (epidermis and trachea). It localises at the apical membrane, where it polymerises chitin polymers that are extruded to the extracellular space. The ability of Kkv to translocate the polymers and deposit them extracellularly depends on the activity exerted by two interchangeable proteins, Expansion (Exp) and Rebuf (Reb). In the absence of Exp/Reb, Kkv can polymerise chitin but it cannot translocate it extracellularly. We recently hypothesised that Exp/Reb regulate the conformational organisation of Kkv at the membrane regulating in this way the translocation activity (De Giorgio et al., 2023). In sharp contrast to Kkv, only little information is available on Chs2. Kkv and Chs2 share common domains, like several transmembrane domains, the conserved catalytic domain QRRRW, and a WGTRE motif, but differ in the presence of a coiled coil domain at the C terminal region in Kkv, which is absent in Chs2 (Moussian, 2013). According to Flybase, Chs2 is mainly expressed in intestinal tissues and is predicted to synthesise chitin, presumably deposited in the PM. A recent paper documented the expression of *Chs2* specifically in a region in the proventriculus in the adult (Zhu et al., 2024) that was proposed to synthesise the Type II PM (King, 1988, Rizki, 1956). However, there is no information on the functional requirements, activity and mechanisms of chitin deposition employed by Chs2.

Here we asked whether Kkv and Chs2 are functionally equivalent and whether they use a comparable mechanism of activity to synthesise chitin. To this end, we have first characterised chitin deposition in the PM and the functional requirements of Chs2 in this process. We have then attempted to replace each CHS with the other one. Finally, we have investigated whether the cellular and molecular mechanisms of chitin deposition by Kkv are shared with Chs2. Our results indicate different specific mechanisms of regulation and activity for each CHS.

## Results

### 1. Chitin accumulation in the intestinal tract

The digestive tract is divided into three main regions: the foregut (which includes the pharynx, esophagus and part of the proventriculus), the midgut (central part of the gut) and the hindgut (starting at the midgut-Malpighian tubules junction) (Miguel-Aliaga et al., 2018), https://flygut.epfl.ch/; (Fig 1A). The digestive tract is lined by a PM that is organised in different layers, L1-L4, visible with electron microscopy approaches (King, 1988). *D. melanogaster* larvae are proposed to produce Type II PM, which is secreted in the proventriculus and slides posteriorly to the midgut enclosing the passing food bolus (Hegedus et al., 2009, Rizki, 1956). Chitin is one of the components of the PM, presumably enriched in the L2 layer of the PM, which helps to organise the glycoprotein mesh that lines the gut (Hegedus et al., 2009, Hegedus et al., 2019, King, 1988, Zhu et al., 2024). We first sought to find a reliable fluorescent probe to visualise chitin “in situ” in the PM.

**Figure 1.**
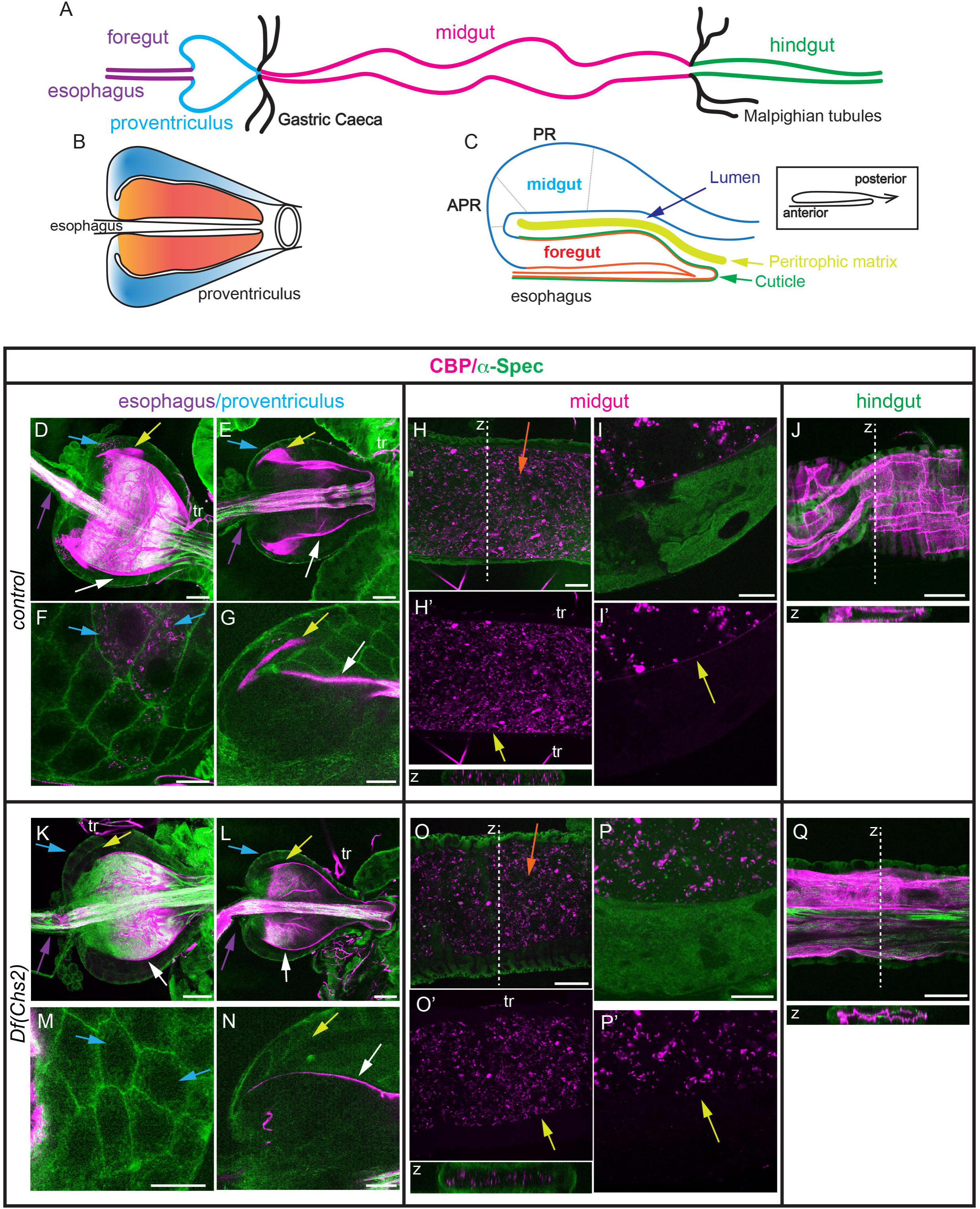
CBP staining along the larval digestive tract. (A-C) Schematics of the larval digestive tract (A), proventriculus (B) and sagital section of half proventriculus (C). The proventriculus forms at the anterior region of the digestive tract and is composed of the esophagus in the inner wall, foregut (ectodermal) cells (orange region in B,C) in the middle wall, and midgut (endodermal) cells (blue region in B,C) in the outer wall. The esophagus and foregut cells are lined by cuticle (green in C). The PM (yellow in C) is deposited in the proventricular lumen and moves posteriorly towards the midgut region (red in A). Different cell types, i.e PR, APR among others, have been identified in the proventriculus region (Rizki, 1956). (D-Q) L3 digestive tracts stained for chitin using CBP (magenta) and Dlg or a-Spec as cellular markers (green). Images show single sections (E-G, J, L-N,Q), projections of 2-3 sections (H,I,O,P) or a projection of several sections (D,K). In the wild type, chitin is deposited in the esophagus (purple arrows), anterior part of the proventriculus (blue arrows), enriched in the proventricular lumen (yellow arrow) and in the PR cells (white arrows) (D-G). In L3 Df(Chs2) escapers the enrichment in the PR region or cells is not detected (K-N). In the midgut region, a continuous layer (yellow arrows) lining the midgut cells is observed in control (H-I’) but not in L3 escapers (O-P’). Note the presence of unspecific CBP staining in the lumen of the gut in control and escapers (orange arrows). No differences are detected in the hindgut (ectodermal) region between control and L3 escapers in chitin deposition (J,Q). Scale bars D,E,H,J,K,L,O,Q 50 μm; F,G,I,M,N,P 20 μm.

A fluorescent-CBP (chitin binding protein) probe, which consists of the chitin binding domain of the Chitinase A1 from *Bacillus circulans,* has been routinely used to visualise chitin in ectodermal tissues (e.g. trachea, epidermis) (Gangishetti et al., 2009). CBP staining, also used “in vivo” (Sobala et al., 2015), nicely and cleanly reveals endogenous and ectopic (intracellular or extracellular) chitin synthesised by Kkv (De Giorgio et al., 2023). We stained the gut of late L3 larvae and detected the presence of CBP in the intestinal tract.

As the PM-chitin is synthesised in the proventriculus (Rizki, 1956), we paid special attention to this region. The proventriculus is formed by the invagination of the esophagus into the midgut, which generates a bulb-shaped structure with 3 layers (King, 1988, Rizki, 1956) (Fig 1B). As described by Rizki (Rizki, 1956), the larval proventriculus has been divided into different regions according to the cell morphology and cell type (Fig 1C, S1A). A group of cells in the outer layer (PR) were proposed to produce the PM. PR cells, as well as cells anterior (APR) and posterior to PR (PPR), correspond to the most anterior midgut cells, while cells anterior to APR correspond to ectodermal foregut cells. We detected a dense, fibrous and strong accumulation of CBP lining the lumen of the esophagus and the lumen of the proventriculus (Fig 1D,E). In addition, the region in the proventriculus that corresponds to PR cells displayed a distinct enrichment of chitin (Fig 1D,E,G, yellow arrow, S1B). We also detected the presence of chitin punctae in the PR cells that seemed to accumulate within the cells (Fig 1F, blue arrow).

Along the midgut region, we detected a continuous, thin and faint layer that lined the gut epithelium (Fig 1H’,I,I’, yellow arrow). In addition, we also detected a homogeneous pattern of CBP staining in the midgut lumen that reveals the accumulation of chitin material along the digestive tract (Fig 1H, orange arrow).

In the most posterior region, the hindgut, we detected a thick and compact CBP staining (Fig 1J).

### 2. Chs2 deposits chitin in the PM

We assumed that the chitin detected in the esophagus and anterior region of the proventriculus corresponds to the cuticle, likely synthesised by Kkv in foregut cells because of its ectodermal origin. In contrast, the chitin enrichment in the lumen of the proventriculus may correspond to chitin deposited in the nascent PM synthesised by PR cells expressing Chs2 because of its endodermal origin ((Rizki, 1956), Fig 1B,C, S1A,B). Consistently, recent data from single-cell RNA sequence analysis indicated that Chs2 is specifically expressed in a corresponding region in the adult proventriculus (Zhu et al., 2024).

To test this, we aimed to analyse the contribution of *Chs2* to chitin deposition in the digestive tract, since no functional analysis of *Chs2* has been performed so far in *D. melanogaster*. We found that a combination of deficiencies that remove Chs2 was not viable, but we could obtain L3 larval escapers (Fig S1C). We found a consistent difference of CBP staining when we compared the escapers to the control. Escapers displayed clear accumulation of chitin in the esophagus and anterior region of the proventriculus, likely corresponding to the ectodermal cuticle, but lacked the enrichment in the PR region (Fig 1K-N, S1C). This suggested that this chitin enrichment in the wild type corresponds to the chitin deposited in the PM by Chs2. We also analysed the pattern of chitin in the midgut. We observed that the faint layer lining the midgut cells was absent in L3 escapers (Fig 1P,P’, S1C). In contrast, we could still detect the homogenous luminal accumulation, indicating that this chitin within the digestive material does not correspond to the chitin synthesised by Chs2 in the PM (Fig 1O). Finally, we detected no differences in the pattern of chitin accumulation in the hindgut (Fig 1Q). We also noticed that the guts of the L3 escapers were thinner and more fragile at dissection.

To confirm that the absence of *Chs2* was responsible for the abnormal pattern of chitin in the *Df(Chs2)* combination, we sought to rescue the phenotype by specific expression of *Chs2*. To this end we generated *Chs2GFP* and *Chs2* transgenes and expressed them specifically in the PR cells by means of a Gal4 line (Zhu et al., 2024) in *a Df(Chs2)* mutant background. Chs2 was able to restore the accumulation of chitin in the PM in the proventriculus and also in the midgut region (Fig 2A-D, S2B-E). Because the *PRGal4* line we used only shows expression in the PR cells ((Fig S2A) and (Zhu et al., 2024)), this result indicates that deposition of chitin in these cells is sufficient to generate the PM-associated chitin lining the whole intestinal tract.

**Figure 2.**
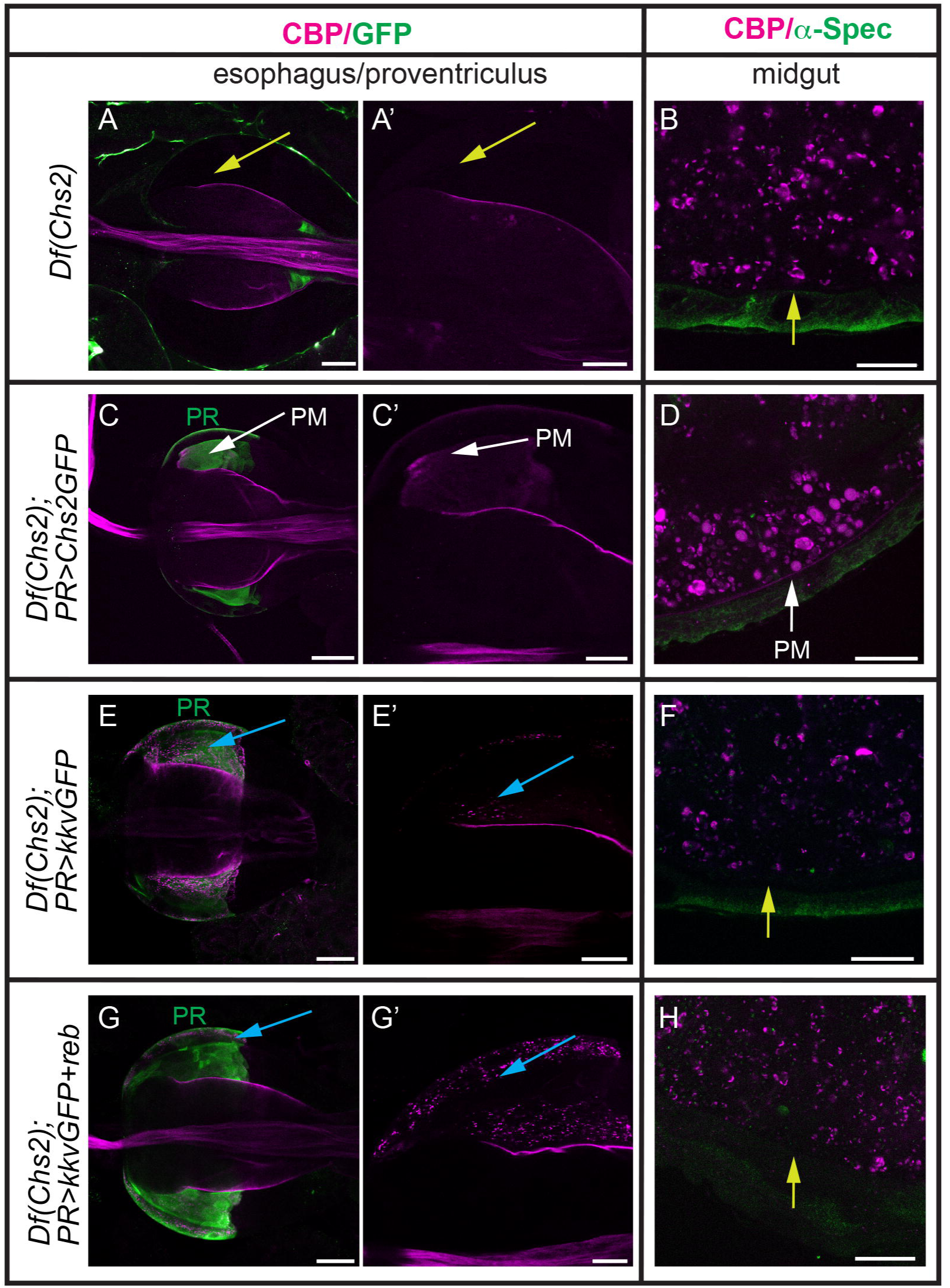
Chs2, but not Kkv, deposits chitin in the PM. (A-C) L3 digestive tracts stained for chitin using CBP (magenta) and α-Spec (A,B,D,F,H) or GFP (C,E,G) in the indicated genotypes. All images correspond to single confocal sections. Note the absence of chitin in the PM in L3 Df(Chs2) escapers (yellow arrows in A,B) and the rescue when Chs2GFP is expressed in PR cells (white arrows in C,D). Expression of KkvGFP alone (E-F) or in combination with reb (G-H) in PR cells results in chitin accumulation in intracellular punctae (blue arrows in E,G) but no chitin deposition in PM (yellow arrows in F,H). Scale bars A,C,E,G 50 μm; A’,B,C’,D,E’,F,G’,H 20 μm.

Altogether we concluded that Chs2 synthesises chitin in the PR cells of the proventriculus. This chitin is deposited in the PM in the narrow proventricular lumen and forms a thin layer that lines the midgut epithelia. In addition, our approach identified a reliable and handy marker to visualise chitin in the PM, CBP.

### 3. Kkv cannot replace Chs2 and deposit chitin in the PM

Kkv deposits chitin in ectodermal tissues (Moussian et al., 2005). We asked whether it could also deposit chitin in the PM. To this purpose we expressed *kkv* in PR cells in a *Df(Chs2)* mutant background. We could not detect chitin in a PM in the proventriculus or in the midgut region. Instead, we detected high amounts of chitin deposited in intracellular punctae in the PR region (Fig 2E,F). This pattern indicated the ability of Kkv to polymerise chitin in the PR region but not to translocate it extracellularly.

We showed previously that chitin translocation absolutely depends on the presence of Exp/Reb (De Giorgio et al., 2023). Thus, we co-expressed *reb* and *kkv* in the PR cells in a *Df(Chs2)* mutant background. We found, again, absence of chitin in the PM in the proventriculus and midgut, and huge amounts of intracellular chitin punctae (Fig 2G,H).

Altogether these results show that Kkv cannot replace the function of Chs2 in depositing PM-chitin even in the presence of Reb.

### 4. Chs2 cannot replace Kkv and deposit chitin in ectodermal tissues

During embryogenesis, Kkv deposits chitin in the epidermal cuticle and in the trachea (Moussian et al., 2005, Moussian et al., 2006). In the trachea, chitin is first deposited inside the lumen forming a filament, and at later stages it is also deposited in the procuticle layer of the tracheal cuticle (Devine et al., 2005, Tonning et al., 2005) (Fig 3A,B). In *kkv* mutants, the tracheal chitin filament does not form and the procuticle layers are devoid of chitin (Devine et al., 2005, Moussian et al., 2015, Moussian et al., 2005, Ostrowski et al., 2002, Tonning et al., 2005) (Fig 3C,D). The expression of *kkv* in tracheal cells of *kkv* mutants perfectly restores the accumulation of chitin in the luminal filament and in the tracheal cuticle (Fig 3E,F) (Moussian et al., 2015). We asked whether Chs2 can substitute for Kkv activity and thus we expressed *Chs2* in the trachea of *kkv* mutants. We found no rescue of chitin accumulation in the filament, in the cuticle or inside the cells (Fig 3G,H, S3A-D).

**Figure 3.**
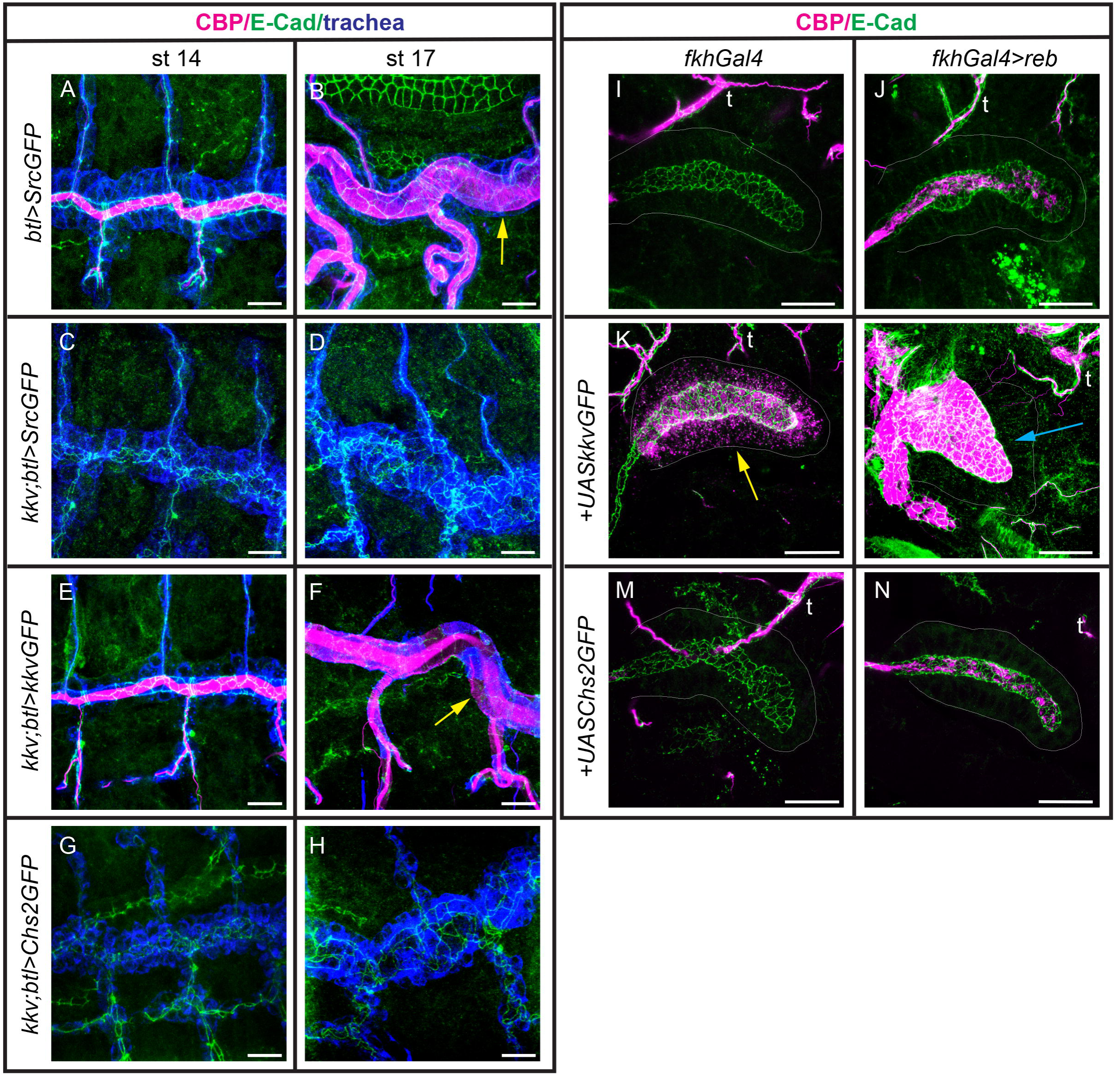
Kkv, but not Chs2, deposits chitin in the trachea and salivary glands. (A-H) Confocal projections showing dorso-lateral views of the trachea stained for chitin (CBP, magenta), GFP (blue) and ECadh (green) to follow the tubes. Note the presence of a luminal filament at early stages (A) and the cuticle with the typical taenidial pattern (B) in the wild type. No chitin in filament or cuticle is deposited in kkv mutants (C,D). Chitin deposition is rescued when adding back kkv (E,F), but not Chs2 (G,H). (I-N) Confocal projections of salivary glands stained for chitin (CBP, magenta) and ECadh (green). Note the capacity of Kkv to polymerise chitin and deposit it intracellularly (K) or in the lumen of the gland when coexpressed with Reb (L). In contrast, Chs2 cannot deposit chitin intracellularly when expressed alone (M). In combination with Reb there is some deposition of chitin in the lumen. This is also observed when Reb is expressed alone (J), and is due to low levels of expression of Kkv in the salivary glands (see Fig S3E). t, trachea Scale bars 10 μm.

We have previously shown that in combination with the activity of Exp/Reb, Kkv is able to polymerise and deposit chitin extracellularly in ectodermal tissues, even in those that normally do not synthesise this polysaccharide (Moussian et al., 2015). In the absence of Exp/Reb, Kkv can polymerise chitin but it cannot translocate and deposit it extracellularly, and as a consequence chitin accumulates intracellularly as membrane-less punctae (De Giorgio et al., 2023). Thus, we observed large amounts of chitin deposited in the lumen when *kkv* and *reb* are co-expressed in the salivary glands (Moussian et al., 2015) and intracellular chitin punctae when only *kkv* is expressed there (Fig 3K,L). The expression of *reb* alone in salivary glands promotes some deposition of chitin in the lumen (Fig 3J) due to the presence of Kkv (Fig S3E,H). We tested the ability of Chs2 to polymerise and deposit chitin when it is expressed in the salivary glands, alone or in combination with Reb. In the absence of Reb, we did not detect accumulation of chitin punctae intracellularly (Fig 3M, S3F). When we co-expressed *Chs2* and *reb* we could not detect increased chitin deposition in the lumen of salivary glands, or intracellularly (Fig 3N, S3G) compared to *reb* expression alone (Fig 3J).

These results indicated that Chs2 cannot replace Kkv function as it is not able to polymerise chitin in ectodermal tissues, even in the presence of Exp/Reb activity.

### 5. Expression of *Chs2* and *kkv* in the proventriculus

To understand better the functional specificity of each CHS we investigated their endogenous pattern of expression. *kkv* is known to be expressed in ectodermal tissues in the embryo (De Giorgio et al., 2023). We investigated its pattern in the proventriculus using a fluorescently tagged knock-in allele (Adler, 2019). We found accumulation of Kkv in the apical domain of the most anterior region of the proventriculus, which corresponds to the ectodermal region, and no accumulation in the endodermal midgut cells of the proventriculus (Fig 4A-C).

**Figure 4.**
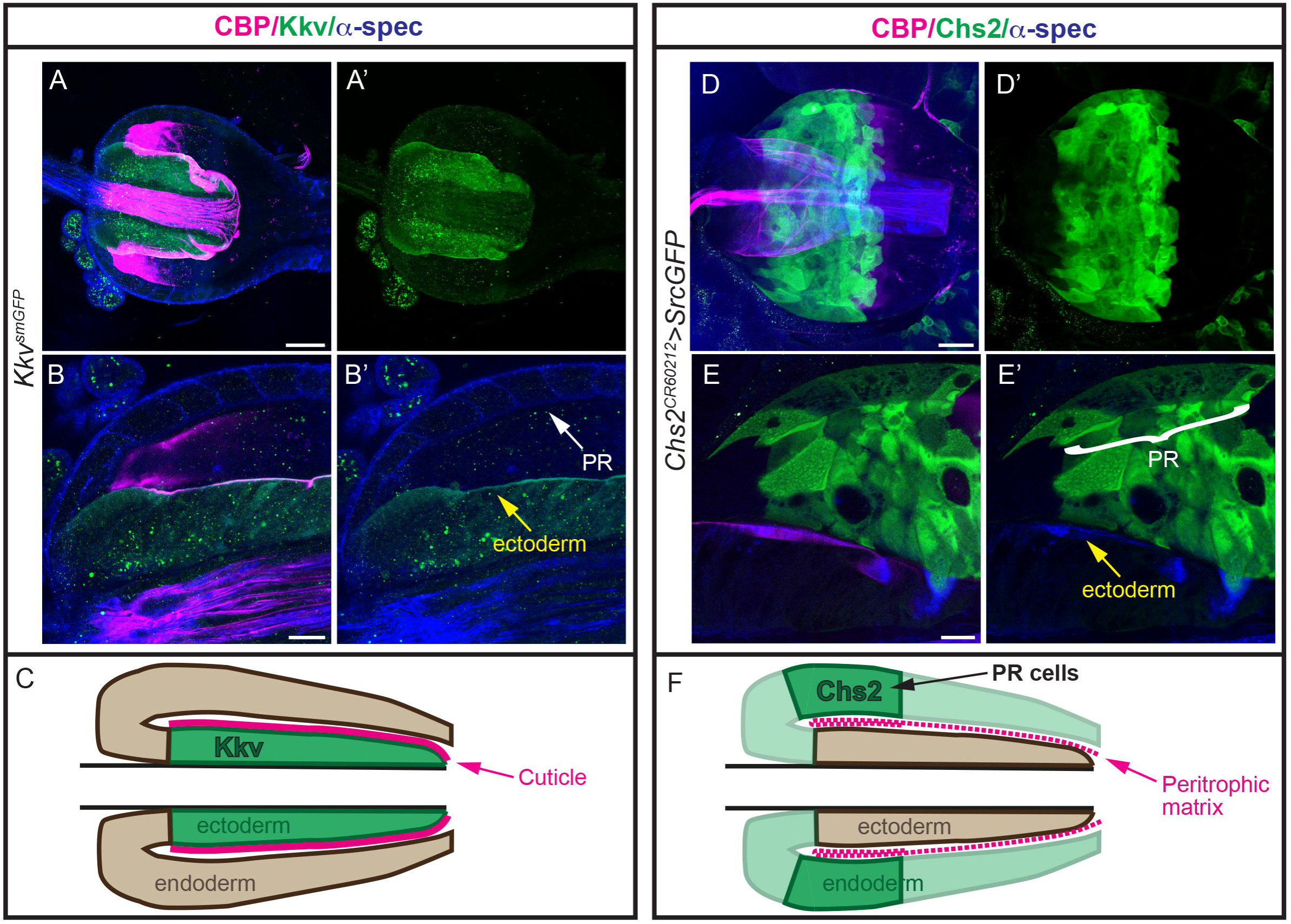
Expression pattern of Kkv and Chs2 in the proventriculus. (A,B,D,E) Single sections or projections of proventriculus stained for chitin (CBP, magenta), α-spec (blue) and GFP (green) to visualise Kkv and and Chs2 pattern. Kkv is expressed in the ectodermal region of the proventriculus (A-C) but not in PR cells. Chs2 is expressed in the PR cells (D-F). Scale bars A,D 50 μm; B,E 20 μm.

A recent report analysing scRNAseq indicated expression of *Chs2* in cells in the proventriculus of the adult that correspond to PR cells (Zhu et al., 2024). To investigate the *in situ* pattern of expression of *Chs2* in the larval proventriculus we used a *Chs2* allele from the CRIMIC collection (Flybase and (Lee et al., 2018)). This revealed clear expression of *Chs2* in PR cells (Fig 4D-F). We also analysed *Chs2* pattern at embryonic stages, and we detected expression in a group of cells at the embryonic proventriculus and an unspecific pattern inside the midgut lumen (Fig S4A,B), but no expression in epidermis or trachea.

Altogether this analysis confirmed the specific expression of Kkv in the ectodermal region and Chs2 in the anterior endodermal region in the proventriculus, consistent with the functional requirements for chitin deposition in the cuticle and PM, respectively.

### 6. Subcellular localisation of Chs2 in endogenous and ectopic tissues

The analysis of the expression of *kkv* and *Chs2* revealed their specific pattern in ectoderm and endoderm respectively. The functional analysis revealed that *kkv* cannot replace *Chs2* activity and that *Chs2* cannot replace *kkv*. To better understand this inability of each CHS to replace the other we analyzed their subcellular localisation.

We first investigated the localisation of Chs2 in the PR cells. We generated an antibody against Chs2 but unfortunately it did not consistently recognise endogenous Chs2 expression in the proventriculus. Thus, we expressed Chs2GFP in PR cells and analysed the localisation. We detected a generalised pattern in the cell but also a clear apical accumulation of the protein that correlates with chitin deposition in the proventricular lumen (Fig 5A-C, S5A). No clear differences in Chs2 localisation were observed in the presence of Reb (Fig S5B). We then expressed Chs2 in other regions in the proventriculus in which Chs2 is not endogenously expressed. Strikingly, while we observed the general pattern in the cell, we could not detect a distinct apical localisation outside the PR region (Fig 5D).

**Figure 5.**
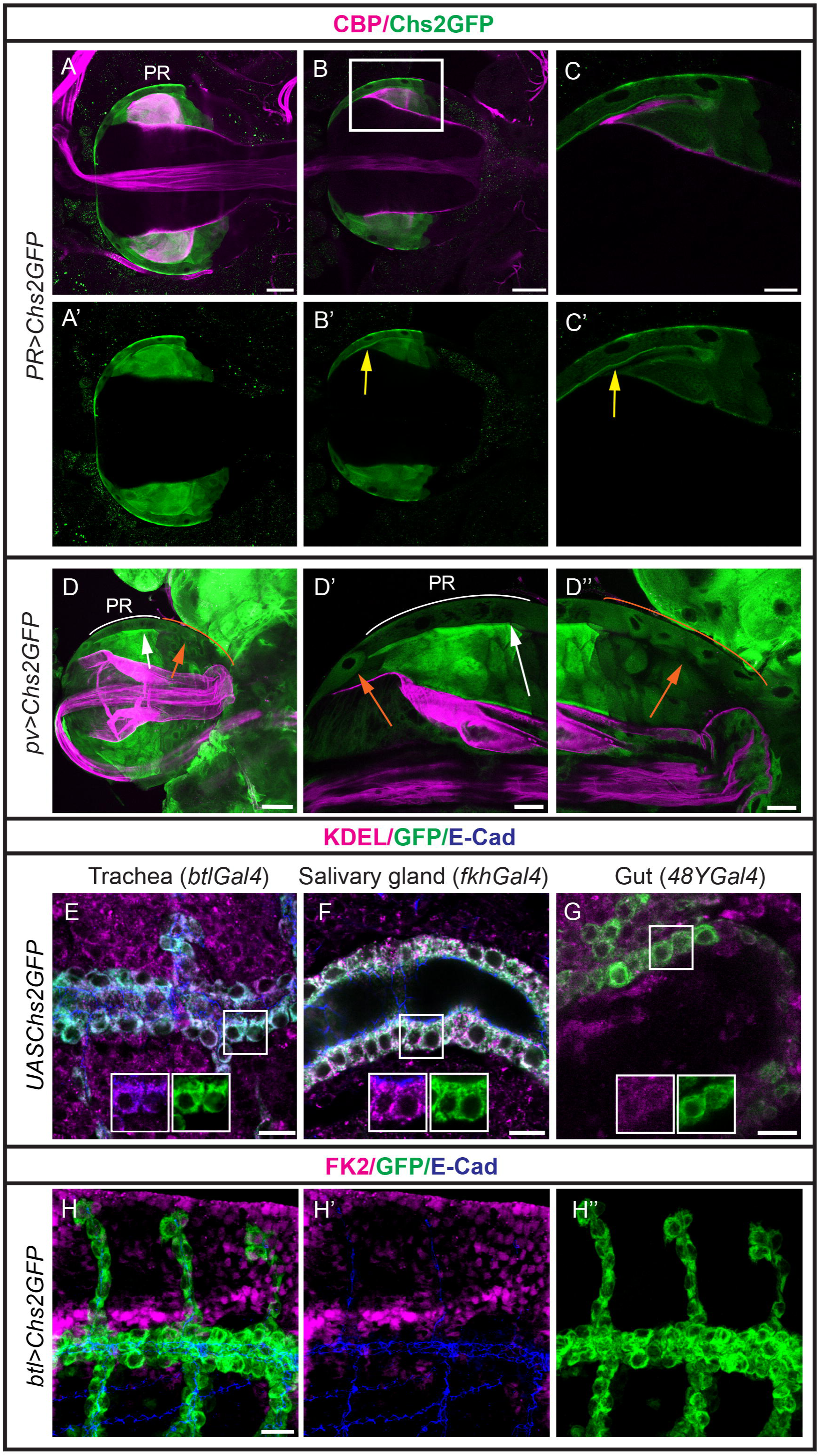
Subcellular localisation of Chs2. (A-D) Proventriculus stained for chitin using CBP (magenta) and GFP (green) in larvae expressing Chs2GFP in the PR cells (A-C) or the whole proventriculus (D). A,D correspond to projections and B,C to single sections of A. C corresponds to a magnification boxed in B. Note the accumulation of Chs2GFP in the apical domain of PR cells (yellow arrows), correlating with chitin deposition. Chs2GFP shows a clear apical localisation only in PR cells (D’, white arch and arrow), in spite of being expressed in the rest of midgut cells of the proventriculus (D’’, orange arch and arrows). D’ and D’’ corresponds to magnifications of D. (E-H) Single sections (E-G) or projections (H) of embryos stained for KDEL or Fk2 (magenta), GFP to visualise Chs2, and ECadh to visualise the cells (blue) in the indicated genotypes. Chs2 localises intracellularly, largely colocalising with ER marker in all embryonic tissues analysed (E-G). Note that Chs2GFP does not colocalise with FK2 (H). Scale bars A,B,D 50 μm; C.D’,D’’ 20 μm, E-H 10 μm

To further analyse Chs2 localisation in ectopic tissues we expressed it in different embryonic tissues. In the ectoderm, like trachea, epidermis and salivary glands, we observed that Chs2 does not localise at the apical membrane. Instead, we detected Chs2 largely accumulated inside the cell (Fig 5E,F, S5D,E), with absent or undetectable accumulation at the membrane (Fig S5C). The cytoplasmic pattern of Chs2 reminded the organisation of the ER, and double staining with KDEL indicated that Chs2 was retained in the ER (Fig 5E,F, S5C-E). We also confirmed that this pattern reflected the localisation of Chs2 protein and was not caused by a missfolding of the protein due to the presence of the GFP in Chs2GFP. We found that the expression of a full-length wild type form of Chs2 in these tissues also indicated ER retention (while the antibody we generated did not recognise clear endogenous pattern of Chs2, it clearly recognised the overexpressed protein) (Fig S5D,E).

We aimed to determine whether Chs2 was specifically retained in the ER or whether the accumulation was due to a defective folding. Missfolded proteins are typically degraded from ER upon ubiquitination (Tsai and Weissman, 2010). FK2 recognises mono and polyubiquitinated conjugates (Fujimuro et al., 1994). We found no colocalisation between FK2 and Chs2 (Fig 5H), strongly suggesting that these proteins are retained in the ER by a specific mechanism/s.

As we have found that Chs2 is endogenously expressed and required in PR midgut cells where it localises apically, we speculated that it could also localise apically in embryonic midgut cells. However, we observed that Chs2 also localised in the cytoplasm in the embryonic midgut, and KDEL co-staining indicated accumulation in the ER (Fig 5G, S5F). This accumulation correlated with the inability of Chs2, alone or in combination with Reb, to deposit chitin in the embryonic gut (a tissue that does not deposit detectable chitin) (Fig S5G,H).

Altogether, our experiments validated the antibody and indicated that Chs2 is retained in the ER when it is expressed in ectopic tissues. In contrast, it localises apically in PR cells in which it is active. These results suggested that a specific factor/s or condition is present in the PR region to promote ER exit of Chs2.

### 7. Subcellular localisation of Kkv in ectopic tissues

We and others have previously shown that Kkv localises at the apical membrane in the tissues in which it is endogenously expressed (Adler, 2019, De Giorgio et al., 2023). In addition, when it is over or ectopically expressed in ectodermal tissues it also localises apically, correlating with its ability to deposit chitin (De Giorgio et al., 2023, Moussian et al., 2015) (Fig 6A,B). We now analysed its localisation when miss-expressed in endodermal tissues. In the embryonic gut we detected a diffuse and general pattern of Kkv in the cell, with some cortical enrichment but no apical localisation. Double staining with KDEL suggested that part of the KkvGFP protein was retained in the endoplasmic reticulum (ER), however, the presence of Kkv containing vesicles suggested that at least part of the protein can exit the ER and traffic within the cell (Fig 6C). This correlated with the ability of Kkv to polymerise chitin intracellularly but not to extrude it extracellularly, even in the presence of Reb (Fig 6D,E). When Kkv was miss-expressed in larval PR cells, in the absence or presence of Reb, also a diffuse pattern was detected. Again, this correlated with accumulation of chitin intracellularly (Fig 6F,G).

**Figure 6.**
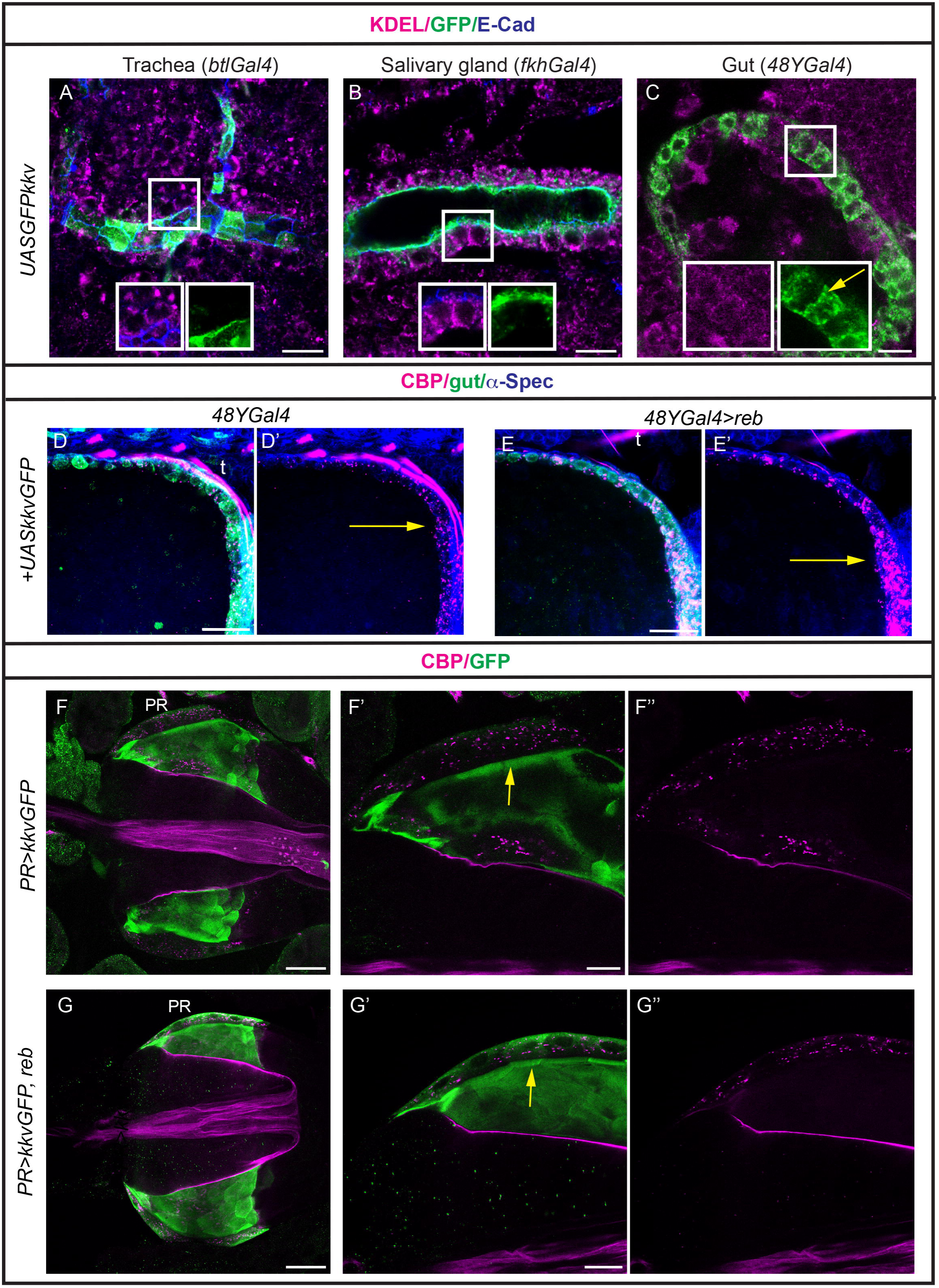
Subcellular localisation of Kkv. (A-C) Single sections of embryos stained for KDEL (magenta), GFP to visualise Kkv and ECadh to visualise the cells (blue) in the indicated genotypes. Kkv localises apically in ectodermal tissues (A,B) and diffusely in the endoderm (C). Note, however, the presence of intracellular vesicles indicating trafficking (yellow arrow in C). (D,E) Single sections of the anterior region of the embryo midgut stained for chitin (CBP, magenta), α-spec as a cellular marker (blue) and GFP (green) to visualise Kkv. kkv can deposit chitin intracellularly in the absence or presence of Reb. (F,G) Single sections of proventriculus stained for chitin (magenta) and GFP (green) to visualise Kkv subcellular localisation in PR cells in the indicated genotypes. Kkv accumulates diffusely in the cell in the absence or presence of Reb (yellow arrows). Clear accumulation in intracellular punctae is detected. t, trachea. Scale bars A-C 10 μm, D-G 20 μm

Altogether this indicated that Kkv can localise at the apical membrane and desposit chitin extracellularly in ectodermal tissues in the presence of Exp/Reb activity. In contrast, it does not localise apically in endodermal derivatives, correlating with its inability to deposit chitin extracellularly, even in the presence of Exp/Reb.

### 8. Analysis of Chs2-Kkv chimeras

At the sequence level, Chs2 lacks a coiled-coil domain that is present in Kkv. We have proposed that this coiled-coil domain plays a role in protein oligomerisation (De Giorgio et al., 2023).

To test the relevance of this domain and determine whether this can explain differences between the two CHS, we decided to generate two different chimeric proteins, one containing Kkv up to the WGTRE domain and Chs2, and a second one containing Chs2 up to WGTRE and Kkv (including the coiled-coil) (kkv-Chs2GFP and Chs2-KkvGFP, respectively, see Fig 7A). We first evaluated their capacity to rescue the absence of chitin in a *kkv* mutant background. We found that none of them was able to restore extracellular chitin deposition or intracellular chitin polymerisation in the trachea of *kkv* mutants, indicating that none of these chimeric proteins are functional to polymerise and deposit chitin in the ectoderm (Fig 7B-D). We then tested whether they were able to replace Chs2, and so we expressed them in PR cells in a *Df(Chs2)* background. We found that none of them were able to deposit chitin in the PM (Fig 7E,F).

**Figure 7.**
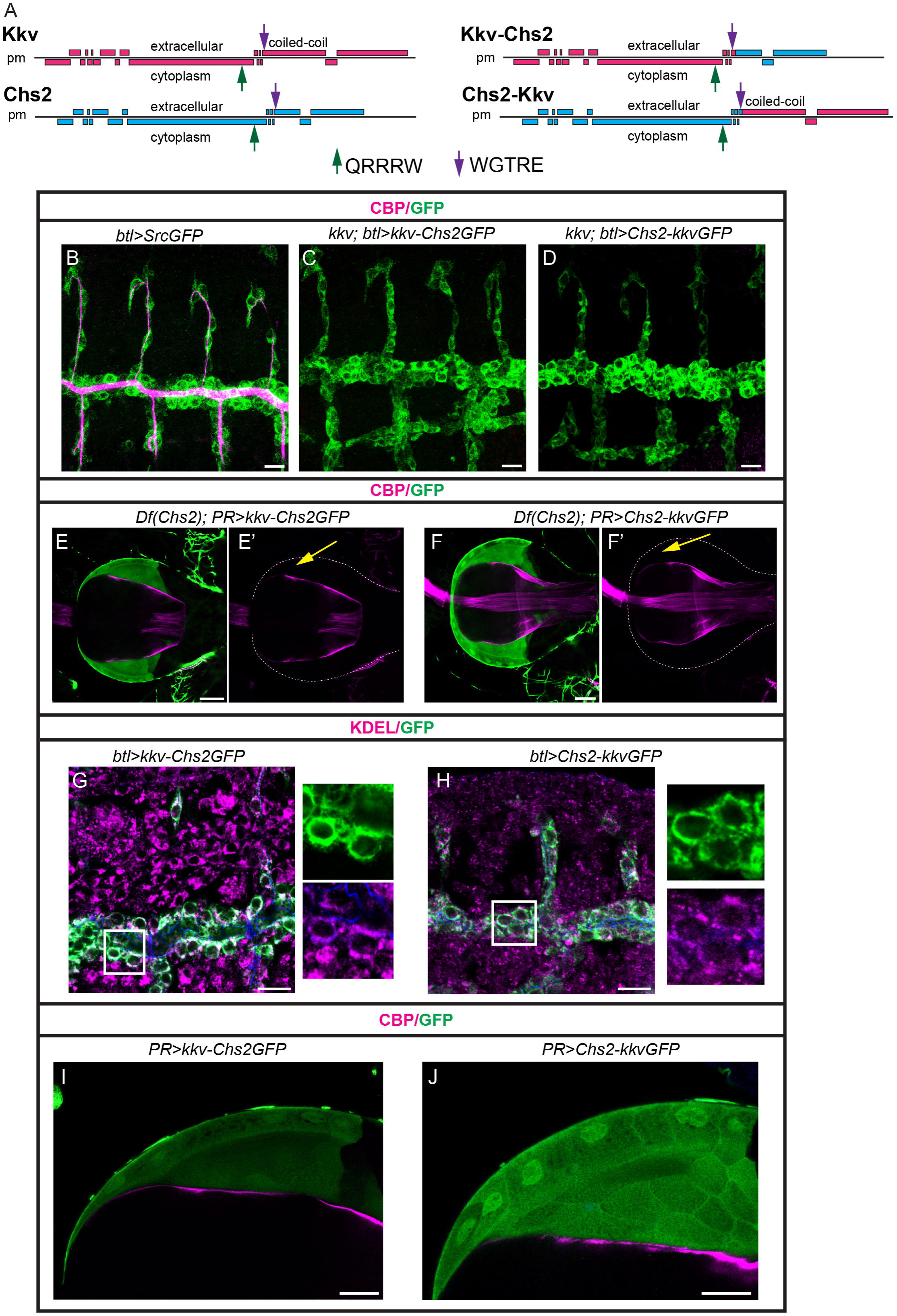
Kkv and Chs2 chimeras. (A) Schematics of Kkv and Chs2 showing the different domains and the position of the catalytic domain (QRRRW) and the WGTRE domain. 2 different chimeric proteins between Kkv and Chs2 were generated as indicated. (B-J) Embryos and proventriculus of the indicated genotypes are shown, stained as indicated. (B-D) The chimeras cannot rescue the absence of chitin in a *kkv* mutant background. (E-F) The chimeras cannot rescue the absence of chitin in the PM in a Chs2 mutant background (yellow arrows point to the absence of PM in the proventricular lumen) (G,H) The chimeras localise intracellularly in embryonic tissues, largely colocalising with the ER marker KDEL. (I,J) The chimeras localise diffusely in PR cells Scale bars B-D,G,H 10 μm; E,F,I,J 20 μm

Finally, we analysed the subcellular accumulation of these chimeric proteins. We found that none of them was able to localise at the membrane in ectodermal tissues, and instead accumulated in the cytoplasm, colocalising with ER markers (Fig 7G,H). In the larval proventriculus, expression in the PR cells indicated a diffuse accumulation within the whole cell with no enrichment at the apical domain (Fig 7I,J)

Altogether our results indicated that the Kkv-Chs2 chimeras are not able to properly localise at the apical membrane and to polymerise and deposit chitin in ectodermal tissues or in PR cells.

## Discussion

In this work we asked about the specificity of chitin synthases in insects, that typically encode two types of CHS, type A and B, with the exception of hemipteran that encode only one type of CHS. While it is well accepted that type A CHS are expressed and required in ectodermal tissues to deposit chitin in cuticles, and type B in the gut to deposit PM-associated chitin, no studies addressed whether they are functionally equivalent. In fact, most functional analyses are based on downregulation of CHS in different insects, addressing only the requirement of their specific tissular expression (reviewed in (Zhao, 2019, Yu et al., 2024)). Here we leveraged the advantages of functional manipulation offered by *D. melanogaster* to ask whether the two different chitin synthases, Kkv and Chs2, are functionally equivalent. We find that a regulated pattern of expression is not the only factor that underlie CHS specificity. In contrast, our analysis indicates that these two chitin synthases are not functionally equivalent and that they use specific cellular and molecular mechanisms to deposit chitin. We suggest that this specificity found for *D. melanogaster* CHS may also apply for the CHS type A and B in other insects.

### Subcellular localisation of chitin synthases

Chitin synthases contain several transmembrane domains and insert in the plasma membrane. They possess a central domain containing the catalytic domain facing the cytoplasm, and a chitin translocating channel with a gate lock through which the nascent chitin filament is extruded (reviewed in (Yu et al., 2024)).

The trafficking of CHS to the plasma membrane has been extensively studied, particularly in yeast. In the well-studied case of *Saccharomyces cerevisiae* Chs3, it is known that the protein is synthesised in the ER, where it folds and oligomerises. Chs3 oligomers then exit the ER assisted by the activity of a dedicated chaperone, Chs7. Chs3 is then transported to the Golgi and is finally delivered to the plasma membrane using the exomer complex. Localisation to the membrane also requires the specific activities of different factors, like Chs4 (reviewed in (Sanchez and Roncero, 2022)). Interestingly, accumulating data indicate that different CHS use different trafficking strategies. For instance, yeast Chs2 is also synthesised and folded in the ER, but it is retained there in a cell-cycle regulated manner: phosphorylated Chs2 is retained in the ER during mitosis due to high mitotic kinase activity, and is exported to the Golgi after mitosis thanks to the dephosphorylation mediated by Cdc14 (Chin et al., 2012, Teh et al., 2009). Thus, besides the need of dedicated proteins and chaperones, different post-translational modifications can play key roles in CHS intracellular transport, also in the case of yeast ScChs3 (Sanchez and Roncero, 2022). In any case, the available data indicate that the regulation of intracellular trafficking and remarkably ER exit represents a critical step for CHS activity, which is conserved in fungi (Rico-Ramirez et al., 2018).

Less is known about the trafficking of CHS in insects, however, several recent advances suggested key roles for intracellular trafficking and ER exit for Kkv functional activity (Chen et al., 2022, De Giorgio et al., 2023, Zhu et al., 2022). In contrast, no previous data existed on *D. melanogaster* Chs2 trafficking. Our studies of Kkv and Chs2 suggest that these different CHS undergo specific transport regulation, and that ER exit is a key regulating step. We find that Kkv is transported to the apical plasma membrane in ectodermal tissues, however, it accumulates throughout the cell in the endoderm. This suggests that either Kkv requires a dedicated chaperone only present in ectodermal tissues or that the intrinsic cell polarity of ectodermal cells provides cues for its apical membrane localisation. We showed previously that the WGTRE domain in Kkv plays a key role in ER exit, suggesting that this domain interacts with a chaperone or undergoes a necessary posttranslational modification (De Giorgio et al., 2023). Chs2 also contains a WGTRE domain, but in contrast to Kkv, an intact WGTRE domain in Chs2 is not sufficient for ER exit in those tissues in which it is not endogenously expressed. This result strongly suggests that Chs2 requires an additional dedicated chaperone or a specific type of posttranslational modification for ER exit and membrane localisation that is only present in PR cells and that is not required by Kkv. In *Manduca sexta* Chs2 localises at the apical brush in the midgut of feeding larvae. In this insect, chitin synthase activity in the gut depends on the regulated expression of Chs2. This regulated Chs2 expression, its apical localisation, and posttranslational modifications that activate the synthase activity, are linked to the nutritional state of the larvae (Broehan et al., 2007, Zimoch et al., 2005, Zimoch and Merzendorfer, 2002). Food ingestion, which starts at larval stages, may also contribute to the regulation of Chs2 activity and PM deposition in *D. melanogaster*. However, the observation that Chs2 localises apically in PR cells and not in the rest of proventriculus cells (which would also interact with the food bolus) strongly points to the presence of a dedicated ER exit factor or posttranslational modification.

Kkv and Chs2 differ in the presence of a coiled-coil domain absent in Chs2. We have recently proposed that this domain plays a role in oligomerisation (De Giorgio et al., 2023). In an attempt to further investigate the role of this coiled-coil domain and determine whether it contributes to the specificity of these two chitin synthases, we designed chimeric proteins in which we added the coiled-coil domain to the Chs2, or we replaced it in Kkv. We found that these two protein chimeras were retained in the ER. We had previously found that the absence of the coiled coil domain in Kkv does not prevent the protein to exit the ER, reach the apical membrane and to be active (De Giorgio et al., 2023). Thus, we speculate that the 2 chimeras we generated are not able to properly fold or oligomerise in the ER, preventing them to reach the membrane. This could indicate that oligomerisation involves both N-terminal and C-terminal domains. Oligomerisation is known to be a critical step in yeast CHS. For example, multiple quality control mechanisms monitor yeast Chs3 folding in the ER, and while non-oligomerised Chs3 can exit the ER, these are returned there once they reach the Golgi using a COPI retrograde trafficking, preventing the trafficking of defective molecules (Sacristan et al., 2013, Sanchez et al., 2023).

### Activity of chitin synthases in *D. melanogaster*

We asked whether Chs2 an Kkv can replace each other. To approach this, we had first to prove that Chs2 deposits chitin in the PM. We were able to establish a reliable and easy method to visualise chitin in the PM by using the widely used chitin probe, CBP. We found chitin deposited at the proventricular lumen and forming a continuous layer lining the whole midgut epithelium. This provides a useful experimental approach for further studies of the PM. We were also able to confirm that Chs2 is the CHS producing PM-associated chitin, as in its absence no chitin was found in the proventricular lumen nor in the midgut. This is a relevant piece of information that was missing in the field. As expected, we found that expressing Chs2 in the PR cells in the proventriculus rescued the lack of PM-associated chitin found in *Df(Chs2)* conditions. Importantly, our results also demonstrated that the expression of Chs2 exclusively in the PR cells is sufficient to produce all chitin deposited in the PM, as it rescued chitin deposition in the proventricular lumen and also in the midgut. This confirms the type II nature of the PM in *D. melanogaster* (Rizki, 1956).

We found that Chs2 cannot replace Kkv and rescue the lack of chitin deposition in the trachea in *kkv* mutants, as Kkv does. Correlating with this observation we find Chs2 retained in the ER when expressed in the trachea. On the other hand, Kkv cannot replace Chs2 when expressed in PR cells, nor even in the presence of Reb. Correlating with this observation we find a generalised pattern of accumulation of Kkv with no apical enrichment, likely suggesting that it does not properly localise at the apical membrane in the brush border region of PR midgut cells (Rizki, 1956). These results indicate specific cellular mechanisms of trafficking for each CHS, that ensure their apical localisation in specific tissues (ectoderm in the case of Kkv and PR cells in the case of Chs2). Insertion of the CHS to the apical membrane represents a necessary requirement for the translocation of the polymers to the extracellular space and therefore for the activity of CHS (De Giorgio et al., 2023, Merzendorfer, 2006, Yu et al., 2024).

Our results also suggest different molecular mechanisms for chitin deposition between Kkv and Chs2. While we find that Kkv requires the function of Exp/Reb (De Giorgio et al., 2023, Moussian et al., 2015), Chs2 seems not to be dependent on this activity. Actually, we observed no differences on the effects of Chs2 expression in the different tissues analysed (trachea, salivary glands, midgut or proventriculus) when Reb is present. scRNAseq detected no expression of *reb/exp* in the midgut cells of the adult proventriculus (Zhu et al., 2024), and we observed no expression of *reb/exp* in PR cells at larval stages. *exp* is expressed in the ectodermal cells of the larval proventriculus (Fig S4C), correlating with *kkv* expression there and cuticle deposition. Thus, we propose that in contrast to the ectodermic Kkv, Chs2 would require a different set of auxiliary proteins and use a specific mechanism to regulate the chitin-translocating channel. Whether this difference results in differences in the structural organisation of chitin promoting specific activities in the cuticles or in the PM is something that remains to be analysed. In this regard, the fact that PM-associated chitin can be visualised by CBP staining provides some information, as CBP recognises insoluble or crystalline chitin but not chito-oligosaccharides or insoluble derivatives of chitin (Hashimoto et al., 2000). While the structural organisation of chitin in cuticles is well studied, it should be noted that there is a lack of structural studies dedicated to understand chitin organisation and its modifications in different insect PM (Merzendorfer, 2016, Hegedus et al., 2009).

In summary, our work shows that that the activity and specificity of *D. melanogaster* CHS relies on different layers of regulation, including the spatio-temporal regulation of their expression pattern, the regulation of their intracellular trafficking, ER exit and membrane insertion, and the organisation of specific molecular machineries for chitin deposition. Our work also provides the grounds and experimental setups for further studies focused on the physiological role of the PM, the specific molecular complexes of chitin deposition used by different CHS, or the structural organisation and properties of chitin fibers produced by different CHS. These are key aspects to understand the full plethora of functional requirements and biotechnological applications of chitin and chitin-modified polymers.

## ACKNOWLEDGEMENTS

We thank L. Portas and C. Rojano for contributions to the initial stages of this project and A. Letizia and ML. Espinàs for continuous help. We thank the IBMB Molecular Imaging Platform for technical help. We acknowledge the Bloomington Stock Centre and the Developmental Studies Hybridoma Bank for fly lines and antibodies.We thank B. Moussian, S. Hayashi and D. Andrew for kindly providing fly lines. We thank the members of the Llimargas and Casanova labs and S. Araújo for helpful discussions, and A. Letizia, ML. Espinàs, J. Casanova and X. Franch-Marro for critical reading of the manuscript.

JB is funded by a FPI Fellowship (Pre2022-102439) and EDG was funded by a FPI Fellowship (BES-2016-076723) of the Spanish Ministerio de Ciencia e Innovación. NM is a technician in Prof. Jordi Casanova’s lab. This work was funded by the Spanish Ministerio de Ciencia e Innovación to Marta Llimargas (MICINN, https://www.ciencia.gob.es/va/), grants numbers PGC2018-098449-B-I00 and PID2021-126689NB-I00). The funders had no role in study design, data collection and analysis, decision to publish, or preparation of the manuscript. The authors declare no competing interests.

## Author contributions

Conceptualization: ML

Methodology: JB, EDG, NM

Validation: JB, EDG, ML

Formal analysis: JB, EDG, ML

Investigation: JB, EDG, ML

Resources: NM

Writing – original draft: ML

Writing –review & editing: JB, EDG, ML

Supervision: ML

Funding acquisition: ML

## MATERIALS AND METHODS

### *D. melanogaster* strains and Maintenance

All *D. melanogaster* strains were raised at 25°C under standard conditions. Balancer chromosomes were used to follow the mutations and constructs of interest in the different chromosomes. For overexpression experiments, we used the Gal4 drivers *btlGal4* (in all tracheal cells, kindly provided by S. Hayashi), *PRGal4* (in PR cells, CG15153^CR02499-TG4.2^, BDSC#92290), *pvGal4* (in all proventriculus, Gnpnat^CR70964-TG4.0^, BDSC#600028), *fkhGal4* (in salivary glands, kindly provided by D. Andrew), and 48YGal4 (in endoderm, BDSC#4935). The overexpression and rescue experiments were performed using the Gal4/UAS system (Brand and Perrimon, 1993).

The following fly strains were used: *kkv^IB22^, UASChs2 and UASKkvGFP* (kindly provided by B. Moussian), and UASsrcGFP. The following stocks were obtained from Bloomington *D. melanogaster* Stock Center (BDSC): y^1^w^1118^ (BDSC#6598, used as control), *Df(3L)BSC233 and Df(3L)BSC284* (BDSC#9700 and BDSC#23669 respectively, deficiencies for *Chs2*), *UAS-reb* (*reb^LA0073^,* BDSC#22192), Chs2^CR60212-TG4.0^ (BDSC#93534), kkv^smGFP-HA^ (BDSC#84993), exp^CR60016-TG4.2^ (BDSC#93301). The UASChs2GFP was generated in our lab.

### Immunohistochemistry

Third instar larvae were dissected in cold PBS and then fixed in 4% Paraformaldehyde for at least 2 hours at room temperature (RT). Larvae were washed 3×15min in PBS-0.3% Triton X-100 (PBT). Dissected guts were blocked with PBT-BSA for 30 minutes and then incubated with the primary antibodies in PBT-BSA overnight at 4°C. Guts were washed with PBS 3×15min and incubated with secondary antibodies in PBT-BSA at RT for 2-4 hours in the dark, washed in PBS and mounted in Vectashield (Vector Laboratories) on microscope glass slides and covered with thin glass slides.

Embryos were stained following standard protocols. Embryos were fixed in 4% formaldehyde (Sigma-Aldrich) in PBS1x-Heptane (1:1) for 10 min for Ecad staining and for 20 min for the rest. Embryos transferred to new tubes were washed in PBT-BSA blocking solution and shaken in a rotator device at room temperature. Embryos were incubated with the primary antibodies in PBT-BSA overnight at 4°C. Secondary antibodies diluted in PBT-BSA (and for the CBP staining) were added after washing and were incubated at room temperature for 2–5 h in the dark. Embryos were washed, mounted on microscope glass slides and covered with thin glass slides.

The following primary antibodies were used: goat anti-GFP (1:600, AbCam); rabbit anti-GFP (1:600, ThermoFisher Scientific – Invitrogen); mouse anti-KDEL (1:200, Stressmarq Biosciences), mouse anti-FK2 (1:50, Enzo Life Science), rat anti-E-Cadh, DCAD2 (1:100, Developmental Studies Hybridoma bank-DSHB, DSHB#528120), mouse anti-α-Spec (1:10 DSHB#528473) and rat anti-Chs2 (1:100, this work).

Cy3-, Cy2- and Cy5-conjugated secondary antibodies (Jackson ImmunoResearch) were used at 1:300. Chitin binding probe fluorescently labelled CBP (1:300) was used to visualise chitin (prepared by N. Martín)

### Image acquisition

Fluorescence confocal images of fixed embryos and digestive tracts were obtained with Leica TCS-SPE system using 20x and 63x (1,40-0,60 oil) objectives (Leica). Fiji (ImageJ) (Schindelin et al., 2012) was used for adjustment. Confocal images are maximum-intensity projections of Z-stack sections or single sections as indicated in the the legends. Figures were assembled with Adobe Illustrator.

### Generation of UAS constructs

Generation of UAS-GFP-Chs2: We amplified the GFP sequence through PCR (Q5 Hot Start High-Fidelity 2X Master Mix, NEB), in the sense primer we included at 5’ the restriction site for XbaI and the Kozak sequence and in the antisense primer at 3’ the restriction sites for NdeI-NheI-XbaI. Upon digestion of the standard cloning vector SK with XbaI, we cloned the amplified sequence of GFP in the plasmid. ChS2 was amplified through PCR and we added at 5’ the restriction site for NdeI and at 3’ the restriction site for NheI. The amplified DNA was cloned into SK-GFP upon digestion of the construct with NdeI/NheI. UAS-GFP-ChS2 (Chs2GFP) was obtained digesting SK-GFP-ChS2 with the restriction enzyme XbaI and cloned in pUAST-attB vector. Upon ligation of the fragments, *E. coli* competent cells were transformed and plated in selective plates. Miniprep to obtain DNA were performed using the kit NZYtech and the DNA was sequenced through the platform Eurofins Genomics. After performing a midiprep (NZYtech), the DNAs were injected in embryos y^1^w^1118^ by the “Drosophila injection Service” of the “Institute for Research in Biomedicine” (IRB, Barcelona) and by the “Transgenesis Service” of the “Centro de Biología Molecular Severo Ochoa” (CBM, Madrid).

The primers used are the following:

PCR primers to amplify GFP: sense 5’-GCT CTA GAG ATG GTG AGC AAG-3’ and antisense 5’-CGT CTA GAA GGC CTG CTA GCC ATA TGC TTG TAC AGC TCC TC-3’;

PCR primers to amplify ChS2: sense 5’-GGA ATT CCA TAT GAG CGG AGG AGAC-3’ and antisense 5’-CTA GCT AGC GCT CTC TGG CGC GTG GGT-3’.

Generation of UAS-GFP-kkv-Chs2 and UAS-GFP-Chs2-Kkv: The constructs UAS-GFP-kkv-Chs2 and UAS-GFP-Chs2-Kkv were obtained by recombination of specific fragments of DNA using the kit “NEBuilder HiFi DNA Assembly” (NEB). The recombination occurs between two sequences of identical DNA, for this reason when we amplified the regions of interest through PCR (Q5 Hot Start High-Fidelity 2X Master Mix, NEB), we added a nucleotide tail complementary to the sequence of the vector and a second tail complementary to the sequence of the other fragment. The DNA templates used during the PCR were pJET1.2-GFP-kkv (De Giorgio et al., 2023) and SK-GFP-ChS2 (described above). We also used the pUAST-attB vector linearised by the restriction enzymes XhoI/KpnI. The kit comprehended material to obtain seamless assembly of multiple DNA fragments. Upon recombination of the fragments, we proceeded as described above with the transformation of *E. coli* competent cells, DNA extraction through miniprep, sequencing of the DNA, DNA extraction through midiprep and, finally, injection of *D. melanogaster* embryos.

The primers used in this study are the following:

PCR primers to amplify GFP-kkv from 5’ until WGTRE motif, with tails overlapping pUAST-attB and Chs2 after WGTRE motif: sense 5’-CTG CGG CCG CGG CTC GAG GGT ACC TCT AGA TGG TGA GCA AGG GCG AGG AG-3’ and antisense 5’-TGA GCA CTG GAG CCT CGC GGG TGC CCC AGGA-3’;

PCR primers to amplify Chs2 after WGTRE motif until 3’, with tails overlapping pUAST-attB and GFP-kkv until WGTRE motif: sense 5’-GGG CAC CCG CGA GGC TCC AGT GCT CAA GGAC-3’ and antisense 5’-AGT AAG GTT CCT TCA CAA AGA TCC TCT AGA ACT CGG TGT GCTC-3’;

PCR primers to amplify GFP-Chs2 from 5’ until WGTRE motif, with tails overlapping pUAST-attB and kkv after WGTRE motif: sense 5’-TCG TTA ACA GAT CTG CGG CCG CGG CTC GAG ATG GTG AGC AAG GGC GAG-3’ and antisense 5’-TCT TAG CCA CCA CCT CTC GAG TGC CCC ACG AAA AC-3’;

PCR primers to amplify kkv after WGTRE motif until 3’, with tails overlapping pUAST-attB and GFP-Chs2 until WGTRE motif: sense 5’-GGG CAC TCG AGA GGT GGT GGC TAA GAA GAC CAA GAA AG-3’ and antisense 5’-CTT CAC AAA GAT CCT CTA GAG GTA CCT CAC AGG CGA CCT GTG CC-3’.

### Generation of antibodies

To generate a polyclonal antibody against Chs2, fragments were amplified by PCR using the following primer combination: sense 5’-CATGCCATGGACTGGTTTCGAACAGGAGGT-3’ and antisense 5’-CCGGAATTCAAAGCCATATTGTTCCG-3’. This amplified a fragment aa F1222 to aa L1383, with low homology to Kkv. The amplified fragments were cloned into the expression vector pPROEXHTa (NcoI/EcoRI). The construct was transformed into Rosetta(DE3) competent cells, and a selected positive clone was induced with 1 mM IPTG at 37°C during 2 hours. The expressed 20 KDa protein fused with a His tag was purified through a column of nickel (Quiagen) in denaturalising conditions (8 M urea). The purified protein was used to inject rabbits by the facility CID-CSIC-Production of antibodies (Barcelona).

---

**Figure S1-Related to Figure 1**

(A) Schematics of a sagital section of half proventriculus indicating the position of the APR,PR and PPR midgut cells

(B) Confocal projection of a wild type proventriculus of a L3 larva stained for chitin (CBP, magenta) and α-Spec as cellular marker (green). PR,APR and PPR cells are indicated. Note the enrichment of chitin in the region of PR cells, corresponding to the PM.

(C) Schematics representing the chromosomal region of Chs2 and the deficiencies used to remove the gene. The combination of the deficiencies gives rise to L3 scapers that lack chitin enrichment in the proventriculus and lining the midgut (yellow arrows), compared to their sibling heterozygote larvae (white arrows point to PM). Scale bars C proventriculus 50 μm; midgut 20 μm

**Figure S2-Related to Figure 2**

(A) Montage of several confocal projections stained for GFP (green) to include the whole digestive tract. Note that the *PRGal4* line used is expressed only in the PR cells and in the hindgut.

(B-E) L3 digestive tracts stained for chitin using CBP (magenta) and α-spec in the indicated genotypes. All images correspond to single confocal sections. No chitin in the PM in L3 Df(Chs2) escapers is observed (yellow arrows in B,C). Expression of Chs2 in PR cells rescues chitin deposition in the PM (white arrows in D,E).

Scale bars A 200 μm; C,D 50 μm; B’,C,D’,E 20 μm.

**Figure S3-Related to Figure 3**

(A-D) Confocal projections showing dorso-lateral views of the trachea stained for chitin (CBP, magenta), GFP to visualise the tracheal cells (blue) and ECadh (green). A luminal filament assembles at early stages (A) and a cuticle with the taenidial pattern forms later (B) in the wild type. Chitin deposition is not rescued rescued in *kkv* mutants when adding back Chs2 (C,D).

(E-H) Single sections (E-G) or projections (E’-G’, H,H’) of salivary glands stained for chitin or Kkv (magenta) and to highlight the salivary glands (green) in the indicated genotypes. Chs2 cannot deposit chitin intracellularly when expressed alone. In combination with Reb there is some deposition of chitin in the lumen (as when expressing Reb alone, Fig 4B’). In *kkv* mutants, the expression of Reb does not promote the deposition of chitin in salivary glands. This indicates that the luminal staining in *fkh>reb* conditions is due to the presence of Kkv. (E,E’) Accordingly, Kkv protein is detected in the salivary glands.

Scale bars A-H 10 μm;

**Figure S4-Related to Figure 4**

Expression pattern of Chs2 in the embryo and Exp in the proventriculus.

(A,B) Single sections of embryos at stage 16 stained with α-spec (magenta) to visualise the cells and GFP to visualise Chs2 pattern. Note the expression in a few cells in the proventriculus. B corresponds to a magnification of A.

(C) Single sections of proventriculus stained for chitin (magenta), GFP (green) to visualise Exp expression and α-spec (blue) to visualise the cells. Exp is expressed in the ectodermal region of the proventriculus and the esophagus and it is absent in PR cells.

ecto-ectoderm; eso-esophagus

Scale bars A,C 50 μm; B,C’ 20 μm

**Figure S5-Related to Figure 5**

(A,B) Single sections of proventriculus stained for chitin (magenta) and GFP (green) to visualise Chs2 subcellular localisation in PR cells in the indicated genotypes. Chs2 is enriched at the apical domain of PR cells (yellow arrows in A’, B’) in the absence or presence of Reb, and its accumulation correlates with a clear enrichment of chitin. Note the fibrous aspect of chitin in the PM (blue arrows in A’’, B’’).

(C-F) Single sections of embryos stained for KDEL (magenta), anti-Chs2 to visualise Chs2 and GFP (blue) to visualise the cell membranes (C) in the indicated genotypes. Chs2 localises intracellularly, largely colocalising with the ER marker in all embryonic tissues analysed (D-F). No accumulation at the membrane level is detected (C).

(G-H) Single sections of the anterior region of the embryo midgut stained for chitin (CBP, magenta), α-spec as a cellular marker (blue) and GFP (green) to visualise Chs2 expressing cells. Chs2 cannot deposit chitin in the embryonic midgut, in the absence or presence of Reb.

t, trachea

Scale bars A,B,G,H 20 μm, C-F 10 μm

