## Supplementary figures and images for "Different cellular and molecular mechanisms of chitin deposition contribute to the specificity of the two chitin synthases in *D. melanogaster*"

### Supplemental Figures

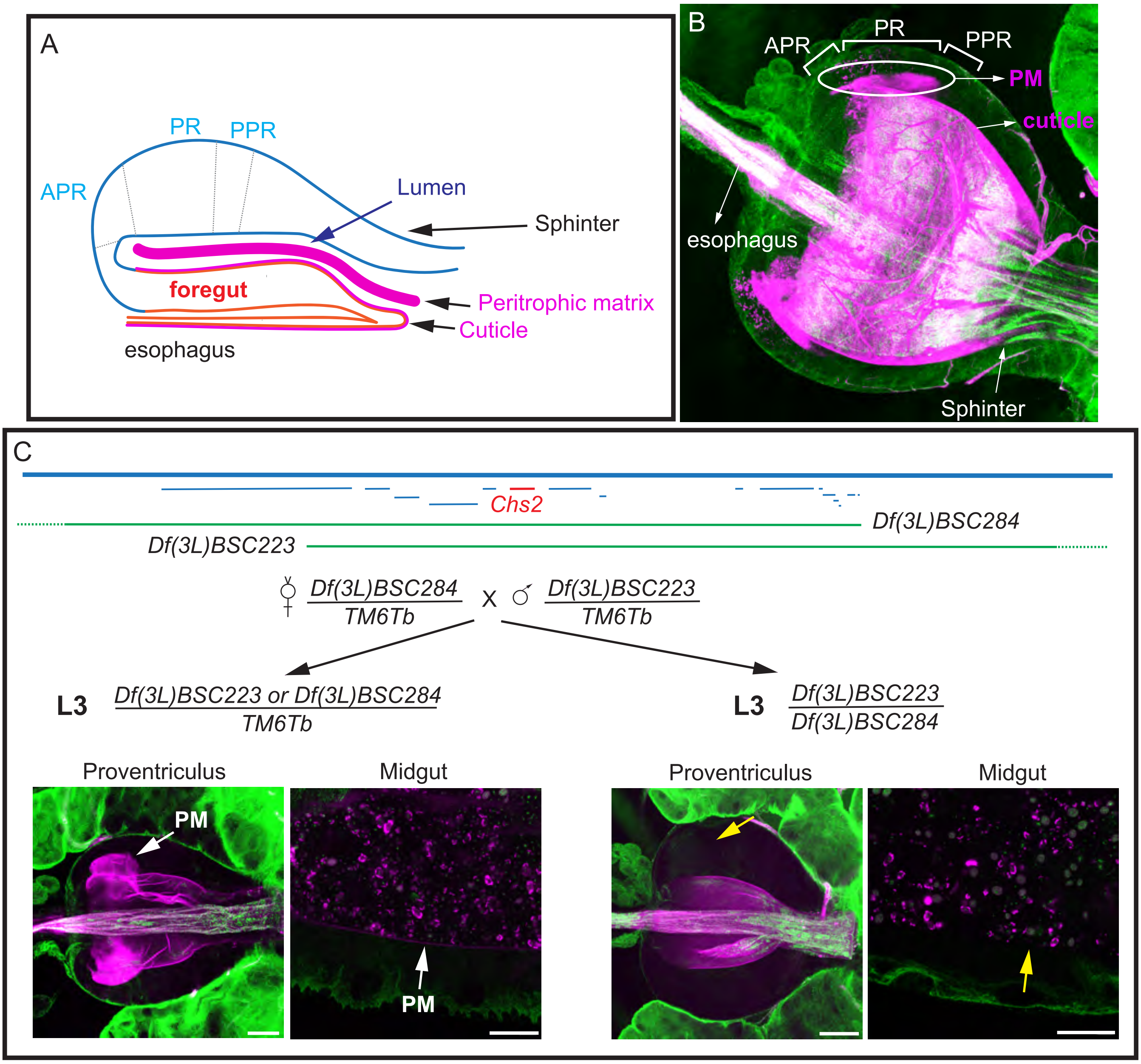

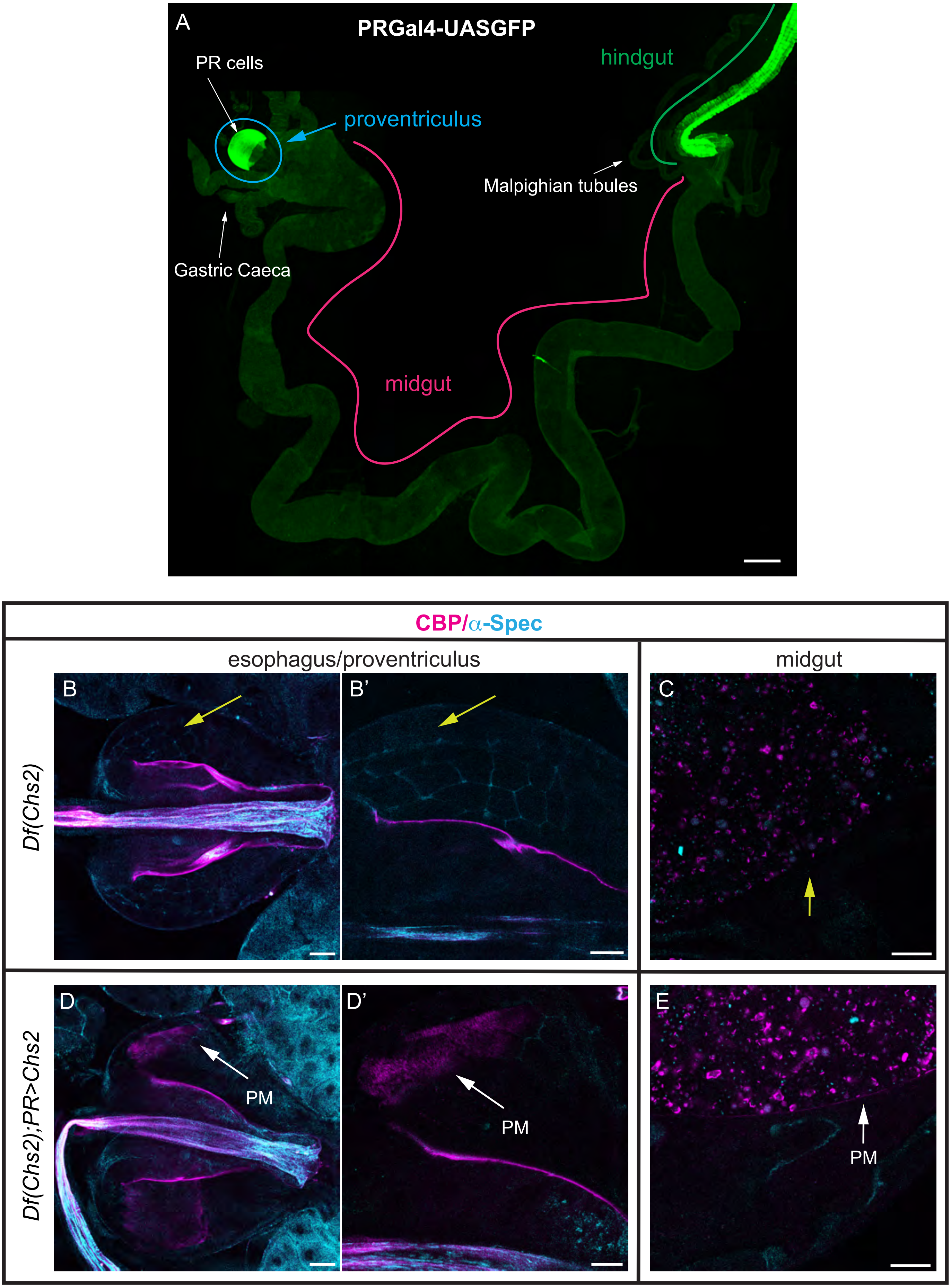

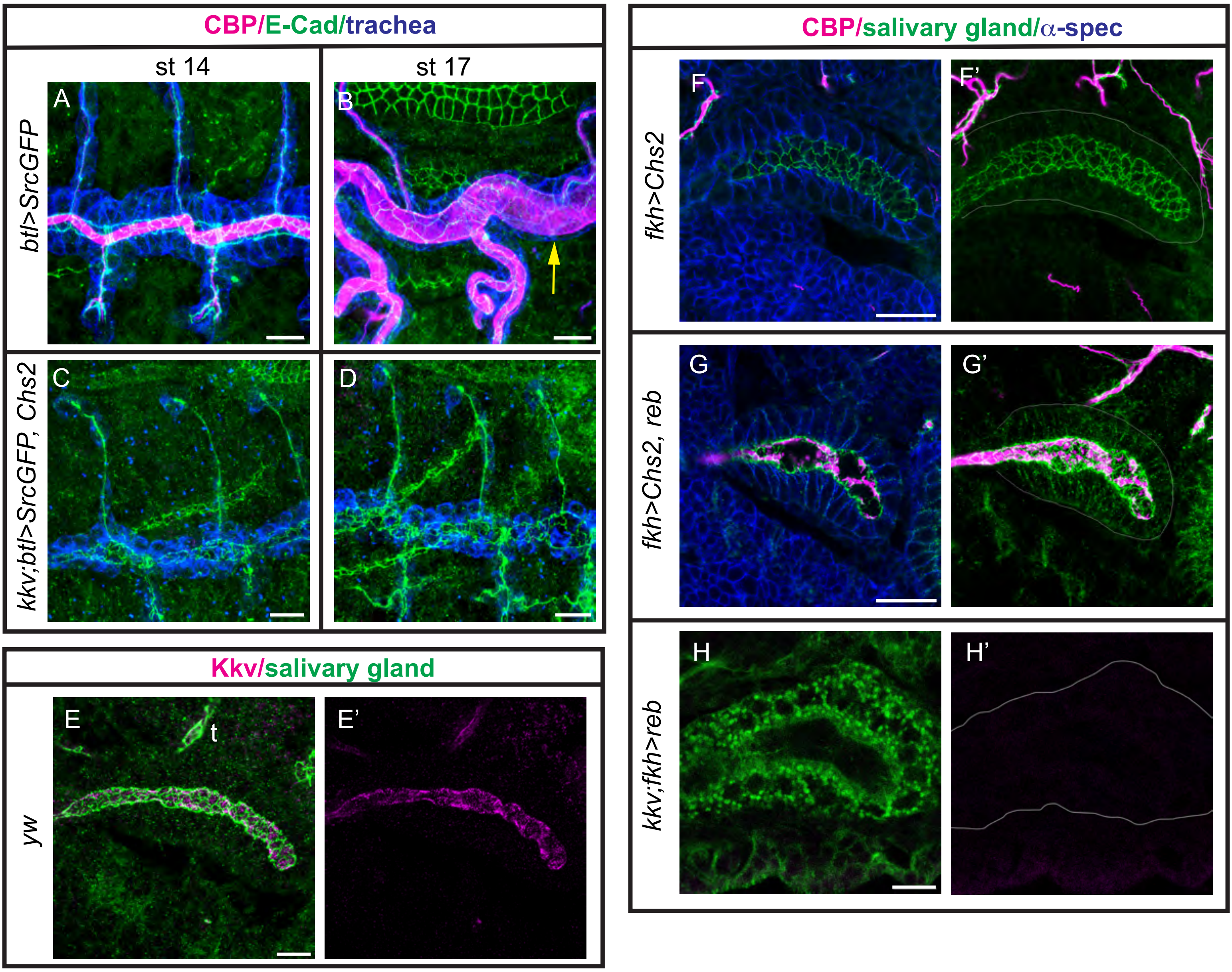

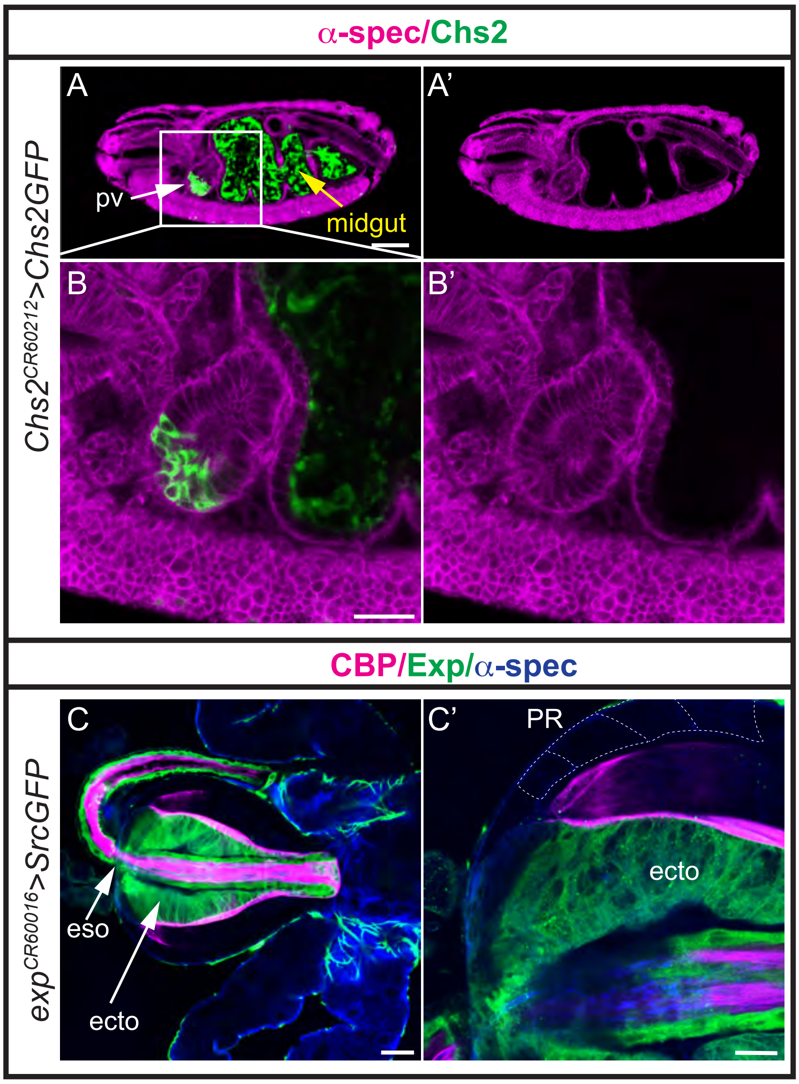

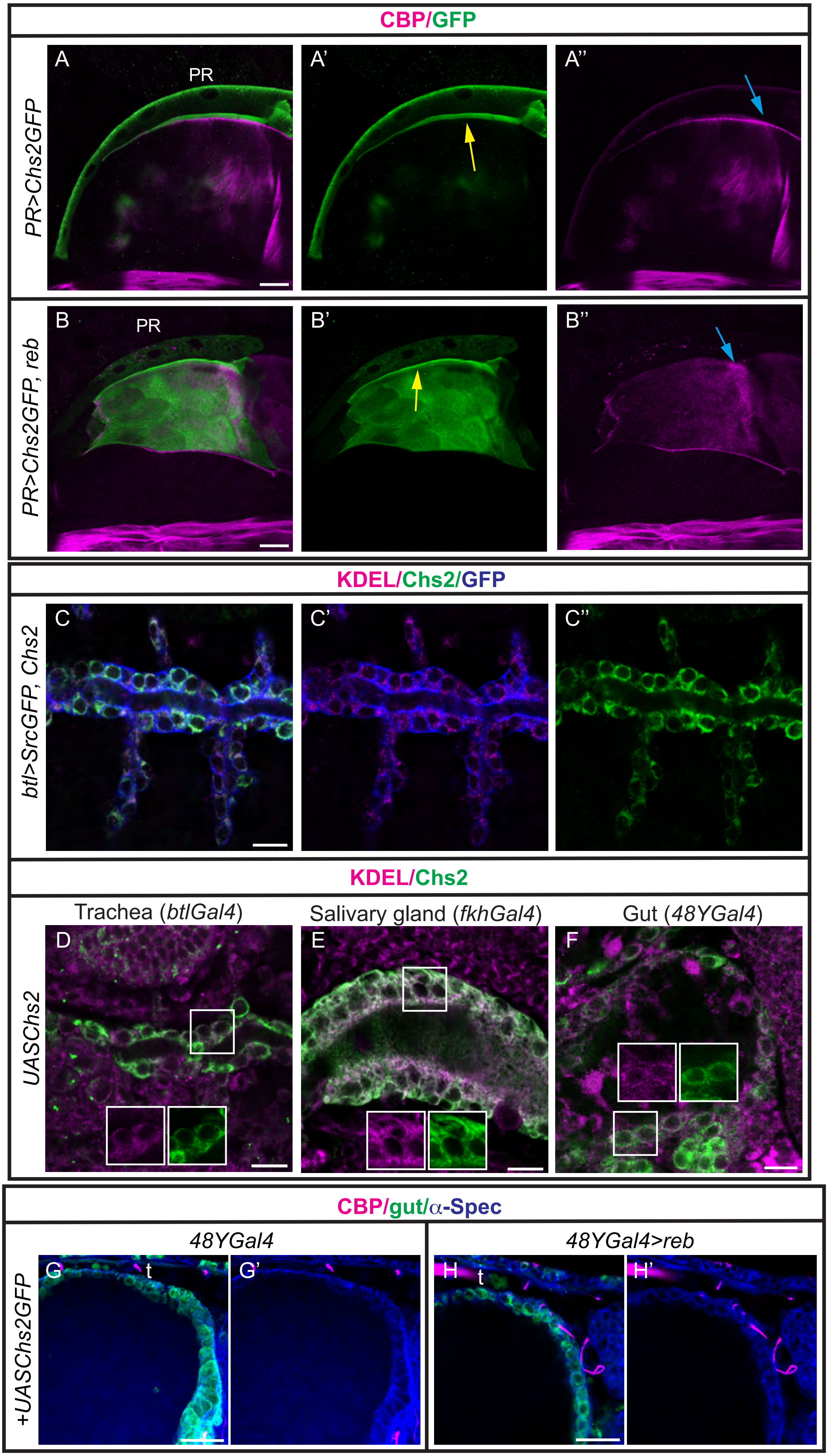
